# Coronin1C SUMOylation modulates filopodia formation, neuritogenesis, and neuronal differentiation

**DOI:** 10.1101/2025.09.29.679180

**Authors:** Garima Joshi, Harsh Vardhan Singh, Ram Kumar Mishra

## Abstract

Actin dynamics in the cytosol and at the cell periphery are critical for metabolic processes and the formation of cellular projections. Coronin1C, a versatile cytoskeletal regulator, is an actin-binding protein that associates with actin at the leading edge of a cell. Consequences of Coronin1C interactions on actin dynamics enable actin-mediated filopodia and neurite formation. Here, we report that Coronin1C is SUMOylated at multiple lysine residues in its carboxy-terminus, preferentially by SUMO1. SUMOylation of Coronin1C is critically required for efficient neurite formation and extensive cellular projections during neuronal differentiation. Moreover, SUMOylation-deficient Coronin1C mutant forms cytoplasmic aggregates colocalizing with stress granules, a hallmark of neurodegenerative diseases, under neuronal differentiation conditions. We also demonstrate that Coronin1C SUMOylation works in synergy with Cdc42 GTPase towards actin-polymerization and promotes filopodia formation. In conclusion, we report that Coronin1C SUMOylation by SUMO1 in its carboxy-terminus is critical for cellular projections formation and neuronal differentiation.

## Introduction

Actin-based filamentous cytoskeletal structures regulate cellular outgrowths, cell migration, vesicular trafficking, and cytokinesis. Filamentous actin (F-actin) pattern and dynamics are regulated by various actin-binding proteins (ABPs), which orchestrate its polymerization and organization. F-actin dynamics is regulated at the base of membrane protrusions such as filopodia, lamellipodia, and neurite formation^1^. One such family of ABPs is Coronins, first identified in *Dictyostelium discoideum* and named after crown-like structures formed on the dorsal surface of these cells^2^*. Dictyostelium* cells carrying Coronin-null mutant exhibit a 50% reduction in cell-migration speed, and are impaired in endocytosis, cytokinesis, and growth^3^.

There are three types of Coronins, and their structural organization is highly conserved, with an amino-terminal domain, a WD40 repeat containing beta barrel domain, and a carboxy-terminal domain. Type I and Type II Coronins share similar structural organization, including an N-terminal extension (N), a *β*-propeller, a C-terminal extension (C), a unique region (U), and a coiled-coil domain (CC). The type I Coronins are major F-actin regulators and are known to coordinate ARP2/3 and cofilin-mediated dynamics of F-actin at the leading edge of the cell^4^.

The 474 amino acid-long Coronin1C (COR1C) is a type I Coronin protein and is involved in regulating cellular migration by aiding in Rac1 trafficking at the membrane protrusions. COR1C depletion in mesenchymal cells decreases the number of cellular protrusions and induces a loss of cellular polarity, thus resulting in impaired migration^5^. The role of COR1C in regulating lamellipodia dynamics has been explored^6^. Multiple studies suggest that elevated levels of COR1C enhance cellular migration and invasiveness, promoting tumorigenesis in different types of cancer^7–12^. COR1C is abundantly present in the brain and is critical for murine brain morphogenesis and migration. Its depletion in Neuro2a cells causes reduced neurite length and number^13^. COR1C binds to actin via its carboxy-terminal coiled-coil domain. The carboxy-terminal domain of COR1C (COR1C-C), and full-length COR1C are indistinguishable as far as subcellular localization and interaction with actin is concerned^14^. COR1C activity is also important for endosome fission^15^. Along with the regulation of actin rearrangement in mouse fibroblasts, COR1C also regulates cytoskeletal dynamics by interacting with vimentin and tubulin^16^.

Post-translational modifications (PTMs) of cytoskeletal proteins are important for maintaining their functions and interactions^17^. Phosphorylation of COR1C at Serine 463 by Casein Kinase 2 modulates ARP2/3 dependent actin bundling and polymerization at the leading edge of the cell^18^. SUMOylation is a reversible attachment of Small Ubiquitin-like Modifier (SUMO) paralogs to target proteins at a conserved lysine residue. It utilizes an enzymatic cascade reminiscent of Ubiquitination. SUMOylation can influence target protein function, stability, and localization^19,20^. A significant number of the cytoskeletal constituent proteins and cytoskeleton-binding proteins are SUMOylated^21^. Interestingly, SUMOylation of nuclear β-actin is indispensable for its nuclear localization^22^. In addition, SUMOylation of Cofilin-1, an actin-filament depolymerizer, affects actin filament dynamics^23^. Coronin family of proteins have been identified as SUMOylation substrates in a global SUMOylation proteomic screen^24^, and Coronin2a is reported to show SIM-mediated interaction with SUMO2/3 ^25^. Can Coronins, particularly COR1C, be SUMOylated, and how does it affect actin-dynamics and cellular processes like cellular migration that are dependent on actin dynamics?

In this study, we report SUMOylation of COR1C at multiple lysine residues (at least five) in its C-terminus by SUMO1. SUMOylation-deficient COR1C-C (COR1C-C-5KR) results in a marked decrease in actin-rich filopodial structures and reduced cellular migration in epithelial cells. Neurite number and length were greatly reduced in Neuro2a cells when SUMOylation-deficient COR1C-C was expressed or when global SUMOylation was inhibited with TAK-981. Interestingly, expression of COR1C-C-5KR under neuronal differentiation conditions in Neuro2a cells induced stress granule colocalizing cytoplasmic aggregate formation that hampered differentiation.

## Results

### Coronin1C is SUMOylated in the Carboxy terminus

Post-translational modification of COR1C by phosphorylation has previously been shown to be critical for its association with ARP2/3 complexes^18^. Moreover, Coronin 2a could interact with SUMO2/3 modified liver-X-receptor through SUMO-interacting motif (SIM)^25^. Proteomic studies identified COR1C to be a SUMOylation-dependent interactor of intermediate filaments (unpublished data). Targeted studies on Coronin family SUMOylation are underexplored, but an extensive proteomic screen reported a smaller isoform of Coronin1C (uniprot ID B4E3S0) to be SUMOylated only at lysine 206 (K206)^24^. We performed *in silico* SUMOylation with prediction tools like GPS-SUMO (SUMOsp.biocuckoo.org) and SUMO JASSA (www.jassa.fr) to predict the SUMOylation sites on canonical 474 amino acid long COR1C (Q9ULV4) (Supplementary Table S1). This analysis revealed that most of the strongly predicted sites (high confidence) lie at the carboxy-terminus of COR1C, referred to as COR1C-C (Fig. 1a) here onwards. To find if COR1C is SUMOylated, we performed *in bacto* SUMOylation assays (see materials and methods for details) with SUMO1 and SUMO2^26^. We found a slow migrating band with anti-GST (for Coronin), and anti-His antibody (for SUMO), indicating SUMOylation with SUMO1 but not with SUMO2 (Fig. 1b). We validated this observation in mammalian cells by co-transfecting GFP-COR1C and Venus-SUMO1 in HEK293T cells and analyzing the cell lysate by immunoblotting using anti-GFP antibody (Fig. 1c). Mimicking *in bacto* observations, a slow migrating band was seen only in the lane representing lysates co-expressing COR1C and SUMO1, thus confirming the COR1C SUMOylation.

**Fig. 1.**
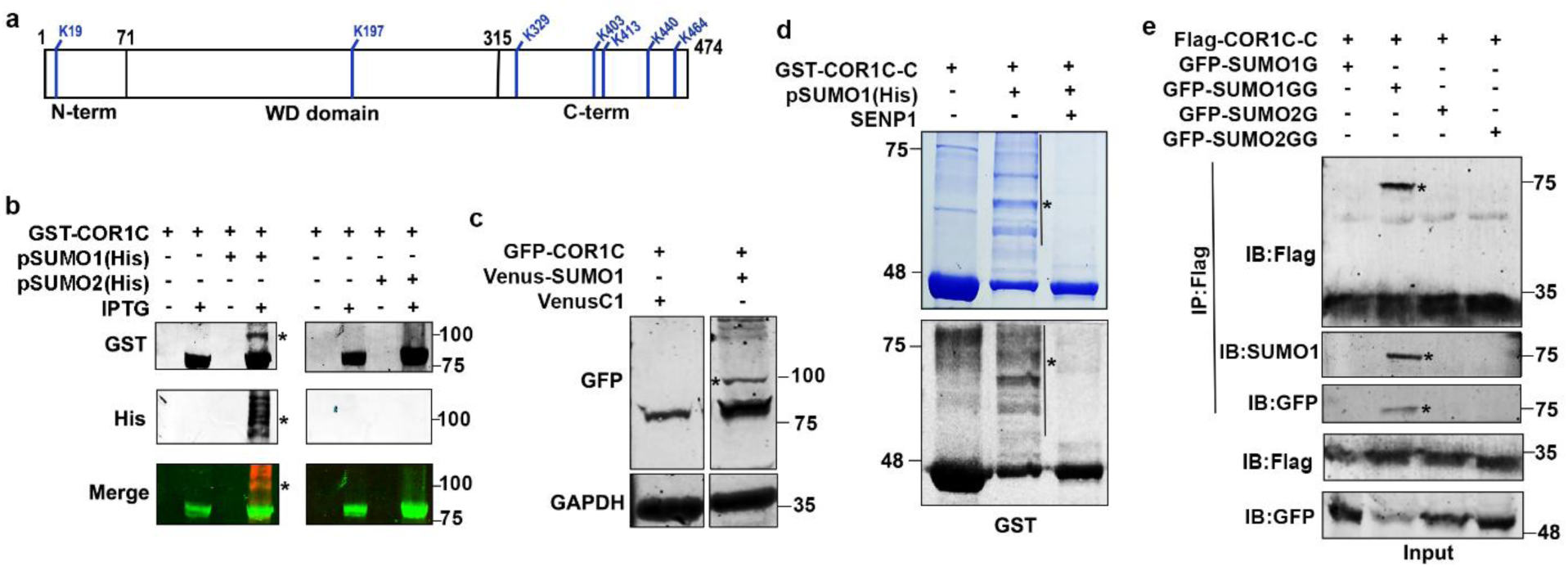
COR1C gets SUMOylated by SUMO1 in its C-terminal domain. **a** Schematic representation of the COR1C protein and its domains. Blue vertical lines indicate lysine residues predicted for SUMOylation. **b** *In bacto* SUMOylation of COR1C with SUMO1 and SUMO2 in the presence of SUMOylation machinery. Modified and unmodified COR1C bands are detected with the anti-GST antibody, and SUMO-modified bands are detected by anti-His antibodies. The bottom panel represents the merged image of both antibodies. **c** Immunoblotting analysis of cell lysates prepared from HEK293-T cells expressing GFP-COR1C alone or in combination with Venus-SUMO1 with anti-GFP antibody. GAPDH is used as a loading control. **d** SDS-PAGE (upper panel) and immunoblotting (anti-GST) analysis of *in bacto* SUMOylation of GST-COR1C-C affinity purified over glutathione beads and treated with SENP1. **e** Lysates prepared from HEK293T cells co-expressing Flag-COR1C-C and GFP-SUMO1G or GFP-SUMO1GG and GFP-SUMO2G or GFP-SUMO2GG were subjected to IP using anti-Flag antibody. IP was analyzed for the presence of any modified COR1C-C bands by WB using anti-Flag, anti-GFP and anti SUMO1 antibody. Input fractions and loading controls are shown. An asterisk indicates a SUMO-modified band.

As most of the predicted sites with high confience lie at the C-terminus of COR1C, we performed *in bacto* SUMOylation with the GST-tagged COR1C-C and found multiple modified bands in Coomassie gel staining (Fig. 1d). Further, we confirmed the *in bacto* SUMOylation of COR1C-C by treating the GST-COR1C-C enriched fractions with the catalytic fragment of SUMO protease, SENP1. A complete disappearance of slow-migrating bands and an increased intensity of COR1C-C bands on SDS-PAGE confirmed COR1C-C SUMOylation (line with asterisk in Fig. 1d, upper panels). A similar observation was made when samples were probed with anti-GST antibody (Fig. 1d, lower panels). Immunoprecipitation of HEK293T cell lysate co-transfected with Flag-COR1C-C and GFP-SUMO1 or SUMO2, revealed a slow migrating band (indicated with an asterisk) with SUMO1 and not with SUMO2 (Fig. 1e). This band was validated to be the modified band by probing with anti-GFP as well as anti-SUMO1 antibody. This experiment confirmed that the C-terminus of COR1C can be SUMOylated in mammalian cells as well.

### The Carboxy-terminus of Coronin1C gets modified at multiple lysine residues

To identify the exact lysine residue(s) on COR1C-C where it gets SUMOylated, the predicted lysine residues (Table S1) were mutated to arginine, individually or in combination with the COR1C-C background, as detailed hereon. The *in bacto* SUMOylation experiments carried out with three individual single lysine mutants, K329R, K403R, and K440R (mutated one at a time), revealed only a subtle, non-significant decrease in SUMOylation when compared with the wild-type COR1C-C. The double mutant K413R, K464R (abbreviated as 2KR) and a triple mutant K329R, K403R, K464R (abbreviated as 3KR) of COR1C-C were probed for SUMOylation. Interestingly, when the extent of SUMOylation for 2KR and 3KR mutants was compared and quantified with the three individual single lysine mutants (described earlier in this section), no appreciable reduction in SUMOylation was noticed (Fig. 2a, b, and Supplementary Fig. S1b).

**Fig. 2.**
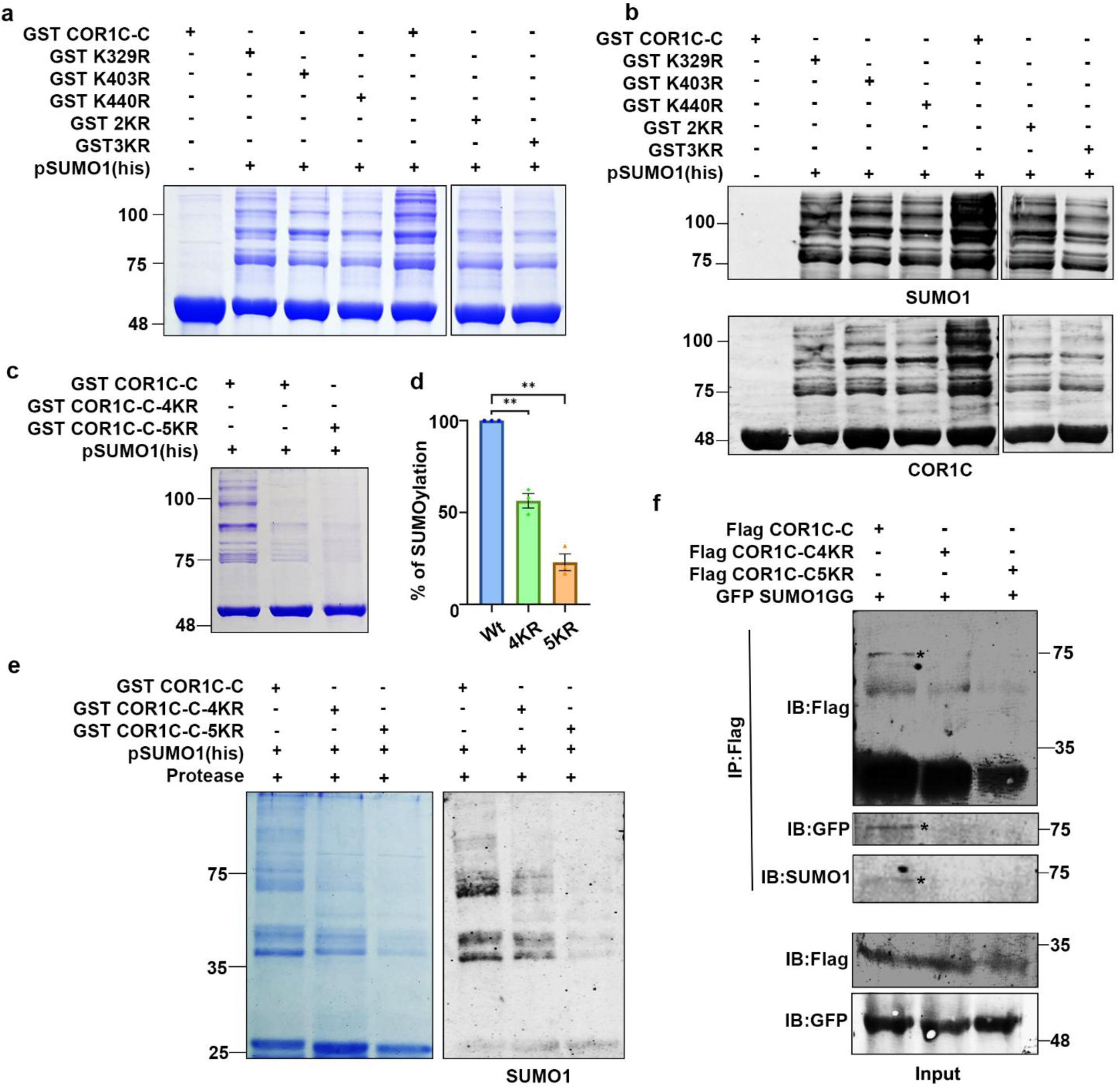
Carboxy terminus of COR1C is SUMOylated at multiple lysine residues.a. Coomassie stained SDS-PAGE analysis of *in bacto* SUMOylation of GST-COR1C-C and its lysine mutants enriched on glutathione affinity beads. **b** Detection of GST-COR1C-C SUMOylation with anti-SUMO1 (upper panel) and anti-COR1C (lower panel). **c** *In bacto* SUMOylation of COR1C-C wild-type and its 4KR and 5KR mutants. **d** Graphical representation of quantification of the percentage of SUMOylation of COR1C-C wild-type and its 4KR and 5KR mutants (n=3 independent experiments). Analysis was performed using an unpaired t-test; (**) indicates statistical significance with a p-value of 0.0081. **e** Coomassie-stained gel and anti-SUMO1 immunoblot of SUMO-modified COR1C-C and its 4KR and 5KR mutants treated with PreScission protease. **f** Lysates prepared from HEK293T cells co-expressing Flag-COR1C-C, Flag-COR1C-C 4KR or Flag-COR1C-C 5KR and GFP-SUMO1GG were subjected to IP using anti-Flag antibody. IP was analyzed for the presence of any modified COR1C-C bands by WB using anti-Flag, anti-GFP and anti-SUMO1 antibodies. Input fractions and loading controls are shown. An asterisk indicates a SUMO-modified band.

There are a total of five predicted lysine residues reflecting high SUMOylation probability. The tested lysines individually or a combination of lysine mutants COR1C-C did not change SUMOylation significantly; hence, we mutated K329, K403, K413, and K464 to arginine to generate COR1C-C 4KR mutant (to be referred to as 4KR from hereon). When the 4KR mutant was probed for *in bacto* SUMOylation and quantified, an average reduction of 56% in the intensity of SUMOylated bands was noticed (Fig. 2c, d). Encouraged by this observation, we mutated 5^th^ lysine also, K440 to arginine, in the COR1C-C-4KR background to generate the COR1C-C-5KR mutant (referred to as 5KR from hereon). Quantification of *in bacto* COR1C-C-5KR SUMOylation reaction SDS-PAGE gels suggested a further decrease to an average of 22.8 % levels (almost ∼75% decrease) as compared to the wild-type COR1C-C modification (Fig. 2c, d). Instead of discrete bands for COR1C-C SUMOylation, many slow-migrating bands were observed, raising a concern that the lysine residues in the GST-tag part of the COR1C-C are also getting SUMOylated. To resolve this, in *bacto* SUMOylation reaction products of COR1C-C wild-type, 4KR, and 5KR mutants were enriched on GSH-beads, and the COR1C-C portion was released from GSH beads using PreScission protease, and analyzed on SDS-PAGE (Fig. 2e, left panel). Here also, we observed slow migrating bands, and probing for their SUMO positivity with anti-SUMO antibody indicated a significant decrease in the 4KR and 5KR mutant SUMOylation, ∼50% and ∼80%, respectively (Fig. 2e, right panel). Further, the immunoprecipitation of COR1C-C mutants from HEK293T cells co-expressing GFP-SUMO1GG and Flag-COR1C-C 4KR or 5KR mutants also confirmed the loss of SUMOylated band in both mutants (Fig. 2f).

### Coronin1C SUMOylation is critical for filopodia formation

Coronins localize at the actin-rich structures of cells, and the involvement of COR1C in lamellipodia formation in fibroblast cells is well documented^27^. More importantly, the primary actin binding sites on COR1C lie in the C-terminus of Coronin1C^28^. Moreover, we know that the expression of COR1C-C colocalizes with F-actin at the cellular projections in HeLa cells (Supplementary Fig. S2) in a manner strikingly similar to the full-length COR1C protein. We next asked if SUMOylation of COR1C-C has an effect on its localization and formation of cellular projections. We overexpressed the GFP-tagged wild-type, 4KR, and 5KR mutant COR1C-C in HeLa cells and quantified filopodial cellular projections. We observed a marked reduction in the number of filopodial projections, 24±2 (for 4KR) and 20±2 (for 5KR), from an average 36±3 (for Wt) filopodial structures per 100 µm cell surface, in cells (Fig. 3a, b). This quantification using FiloQuant^29^ indicated ∼ 33-50% reduction in filopodia number in the mutants. Similar observations regarding reduction in the filopodia numbers upon GFP-COR1C-C mutant expression were made from MCF7 cells as well, confirming the importance of COR1C SUMOylation in a cell type-independent manner (Supplementary Fig. S3). Interestingly, the average length of a filopodia remained unchanged in cells expressing the GFP-COR1C-C-4KR or 5KR mutants (Supplementary Fig. S4). To further confirm the impact of SUMOylation on filopodia formation, we treated GFP-COR1C-C expressing HeLa cells with TAK981, a potent inhibitor of E1 enzyme of the SUMOylation pathway, to cease global SUMOylation. Quantification from TAK981 treated HeLa cells indicated an approximately two-fold reduction (from 39±4 in control to 15±1 in TAK981 treated) in the formation of filopodial projections on these cell surfaces, an observation strikingly similar to what is seen with the COR1C-C-5KR mutant (Fig. 3c, d).

**Fig. 3.**
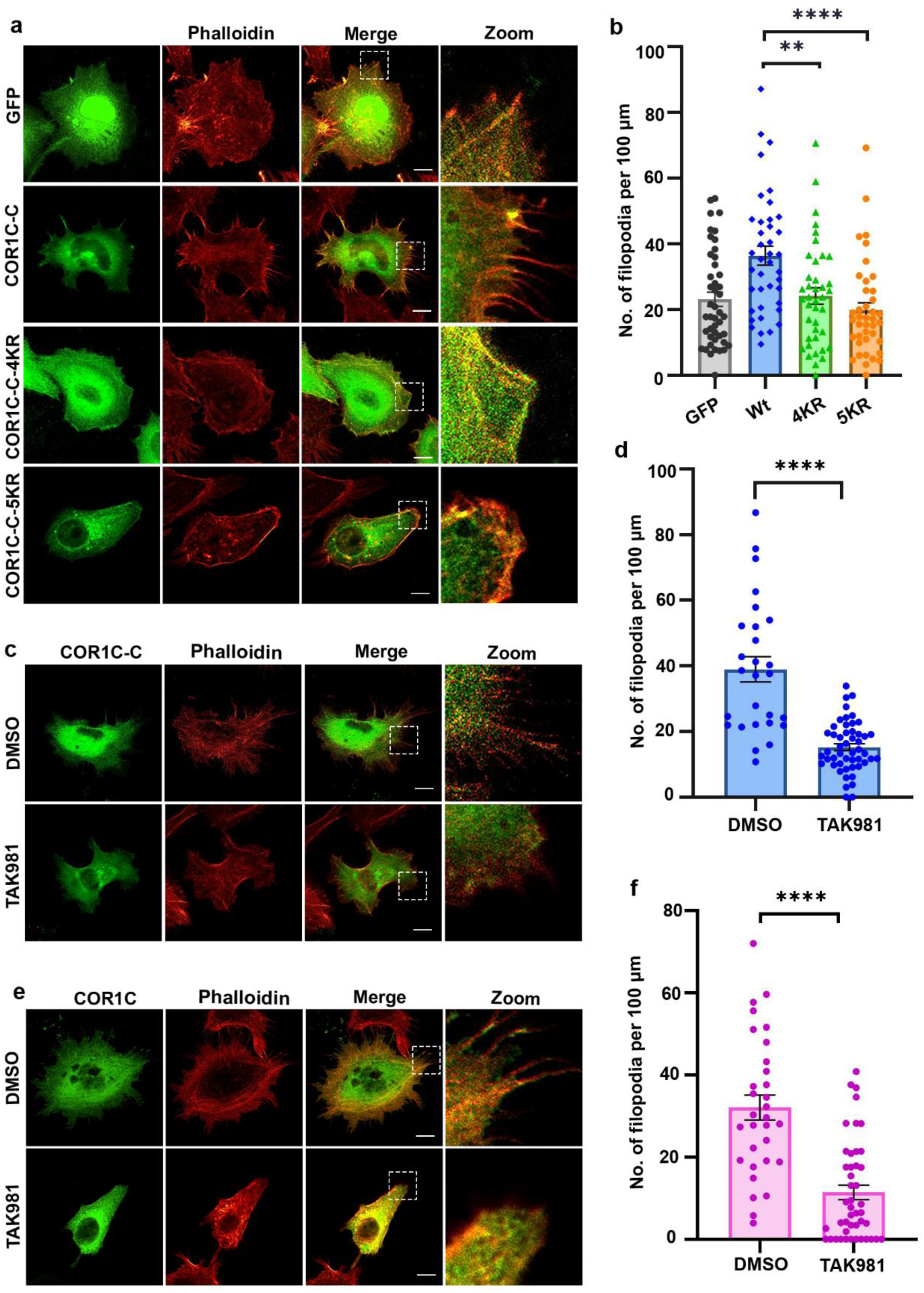
COR1C SUMOylation is critical for filopodia formation in cells. **a** CLSM imaging of HeLa cells transfected with GFP, GFP tagged COR1C-C, COR1C-C-4KR, COR1C-C-5KR. Cells were stained with Phalloidin-647. The dotted box highlights filopodial structures in the leading edges, as shown in the zoom. **b** Filopodial projections in (a) were quantified using the FiloQuant plugin of ImageJ (∼ 50 cells were analyzed in 3 independent experiments). Asterisks (**) and (****) indicate statistical significance with a p-value of 0.0017 and <0.0001, respectively. CLSM imaging of GFP-COR1C-C (**c**) and GFP-COR1C (**e**) expressing HeLa cells treated with DMSO or 5 μM TAK981 for 48 hours. **d, f** Filopodial projections in (c) and (e) were quantified using the FiloQuant plugin of ImageJ (∼ 50 cells were analyzed in 3 independent experiments). (****) indicate statistical significance with p<0.0001, respectively. Statistical analysis was performed using an unpaired t-test. Scale bar – 10 μm.

Similarly, quantification from full-length GFP-COR1C expressing HeLa cells, treated with TAK981, shows a reduction of > 2-fold in filopodia numbers from 32±3 in control to 11±2 in TAK-981 treated cells (Fig. 3e, f). This suggests that SUMOylation of COR1C is important in its role in filopodia formation.

A reduction in filopodia numbers can manifest an obvious migrating defect in adherent cells^30^. We performed a wound healing assay with wild-type GFP-COR1C-C or the 5KR mutant expressing HeLa cells (as described in methods) to assess if filopodia formation dependent on COR1C-C SUMOylation can alter cell migration. Cells expressing COR1C-C wild-type migrated faster than either GFP or 5KR mutant expressing cells, closing the wound almost completely in 48 hours. The area of wound closure in GFP expressing cells was 86.3%, whereas in the COR1C-C expressing cells it increased up to 93.7%. In the 5KR mutant expressing cells, the wound closure was 85.8% which was closer to the GFP control, thus suggesting that the overexpressing COR1C-C in HeLa cells allow them to migrate faster. The delay in wound closure observed in 5KR mutant conditions highlighted the importance of COR1C-C SUMOylation in filopodia formation and cell migration (Supplementary Fig. S5a, b). We argue that the migration delay under 5KR mutant expression conditions could be more pronounced. But due to the presence of endogenous COR1C protein in cells transiently expressing 5KR mutant, wound closure defects are less pronounced.

### COR1C SUMOylation works synergistically with Cdc42-dependent actin dynamics during filopodia formation

Generally, filopodia formation is an actin dynamics-dependent process, mostly governed by Cdc42 GTPase^31^. Coronin 1C is an actin-binding protein. We hypothesize that the SUMOylation of COR1C-C is required for regulating actin polymerization dynamics during the formation of cellular processes.

ML141 is a Cdc42 specific inhibitor, routinely used to inhibit filopodia formation^32^. In order to check the Cdc42-dependence of filopodia formation impairment in the cells expressing COR1C-C 5KR mutant, we treated the GFP, COR1C-C and COR1C-C 5KR mutant expressing cells with ML141 and observed filopodia formation. The ML141 treated cells expressing GFP or GFP-COR1C-C-5KR mutants show significant defects in the number of filopodia formed. In contrast, the GFP-COR1C-C expressing cells resisted this decline and observed a modest decrease in filopodia number (Fig. 4a). The quantification of this data suggested a decrease of 53% (from 20±2 to 8±3 per 100 μm) and 45.6% (from (23±2 to 13±2 per 100 μm) in filopodia number in ML141 treated 5KR mutant and GFP expressing cells, respectively. Whereas this decrease in the filopodia number was found to be only 28.5% (from 36±3 to 26±3 per 100 μm) in COR1C-C overexpressing cells (Fig. 4 b). In addition to this, we also observed a stark decrease in the length of filopodia in ML141 treated cells in all the conditions (Fig 4c). The length decreased from 0.85±0.07 to 0.22±0.023 μm in GFP expressing cells, from 0.96±0.05 to 0.33±0.023 in COR1C-C expressing cells, and from 0.83±0.06 μm to 0.21±0.04 μm in 5KR expressing cells. It is interesting to note that the reduction in length was more drastic in the case of 5KR mutants when compared to the wild type when Cdc42 was inhibited, but under normal conditions, the filopodia length remained unperturbed, as previously shown in Supplementary Fig. S4.

**Fig. 4.**
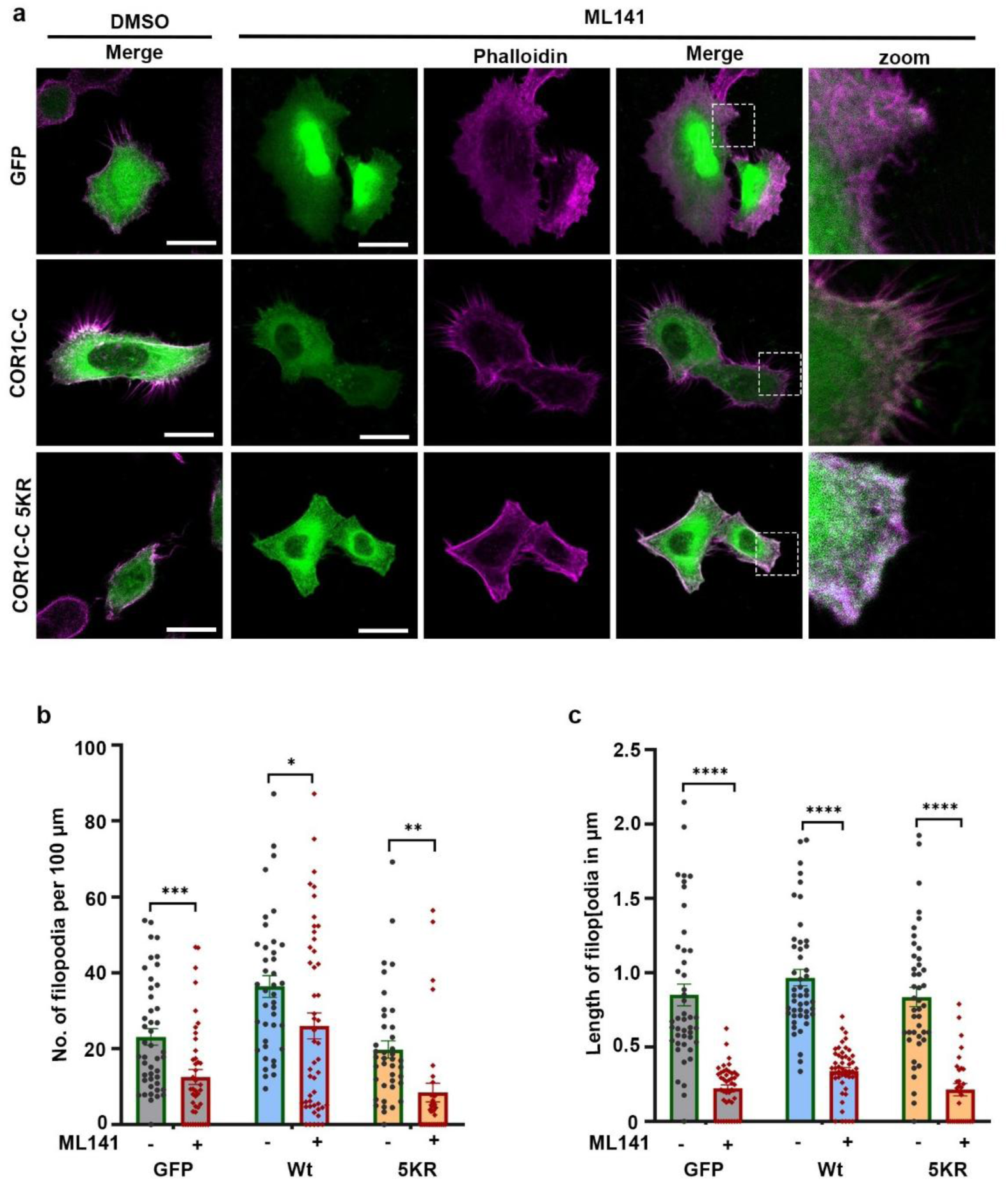
COR1C SUMOylation and Cdc42 work synergistically during filopodia formation. **a**. HeLa cells were transfected with GFP, GFP-tagged COR1C-C, and COR1C-C-5KR and imaged after 24 hours using a CLSM after a treatment with ML141(10 μM) for 2 hours. F-actin was stained with Phalloidin-647. Scale bar - 10 μm. **b.** Filopodial projections in (a) were quantified using the FiloQuant plugin of ImageJ (at least 35 cells were analyzed in 3 independent experiments). Asterisks (*), (**), and (***) indicate statistical significance with a p-value of 0.0253, 0.0049, and 0.0005, respectively. **c** Quantification of the length of filopodial projections measured from HeLa cells (a) using FiloQuant plugin of ImageJ.

This strongly suggests that COR1C SUMOylation and Cdc42 work synergistically during filopodia formation. Perturbing COR1C SUMOylation or inhibiting Cdc42 activity, both compromise filopodia formation. Our observations suggest that COR1C promotes filopodia formation at cellular edges, and its SUMOylation plays a critical role in this process.

### SUMOylation of Coronin1C enhances neuritogenesis in Neuro2a cells

Neuritogenesis is a process during neuronal differentiation where neurons form axons and dendrites^33^. In neurons, filopodia-like projections dilate and mature to form neurites during neuronal development and differentiation^34^. We have observed that COR1C SUMOylation is important for filopodia formation; therefore, we asked if COR1C SUMOylation can also impact neurite formation and neuronal differentiation. In this context, we have quantified multiple neurite-related aspects like the presence of neurites, their length, and the number of neurites per cell in Neuro2a (N2A) cells. Neurite-related attributes were observed and quantified from GFP, GFP-COR1C-C, GFP-COR1C-C-4KR, or GFP-COR1C-C-5KR expressing N2A cells. Normally, the cell body size of N2A cells is 15-20 µm, and in our calculations, we consider cellular projections longer than 50 % of the cell body size as neurites. The percentage of cells positive for neurite increased by almost two-fold in GFP-COR1C-C expressing cells, when compared with GFP expressing control cells (Fig. 5a, b). In contrast, in the cells expressing GFP-COR1C-C-4KR and GFP-COR1C-C-5KR mutants, the number of neurite-positive cells remained unchanged when compared with GFP-expressing control cells (Fig. 5a, b). Careful analysis of neurite length indicated that the length of neurites produced in COR1C-C-4KR or 5KR mutant expressing cells is significantly reduced (from 17.3±0.7 µm in wild-type to 13.62±0.54 µm for 4KR and 11.9±0.63 µm for 5KR mutant) (Fig. 5c). The number of neurites per cell is an indication of neuronal development and differentiation, so we counted the number of neurites per cell. Cells with no neurites decreased to ∼ 12% in GFP-COR1C-C expressing cells from ∼ 48% in GFP expressing control cells. Moreover, the number of cells having more than 3 neurites per cell increased to ∼ 25% in COR1C-C expressing cells from ∼ 5% in GFP expressing cells. More importantly, the percentage of cells with no neurites and cells with more than 3 neurites in COR1C-4KR or 5KR mutant expressing cells remained comparable to GFP expressing cells (Fig. 5d). Together, these data suggested that COR1C-C expression induces neuritogenesis and associated key aspects of differentiation, and COR1C-C SUMOylation is integral to COR1C-mediated neuritogenesis and neuronal differentiation.

**Fig. 5.**
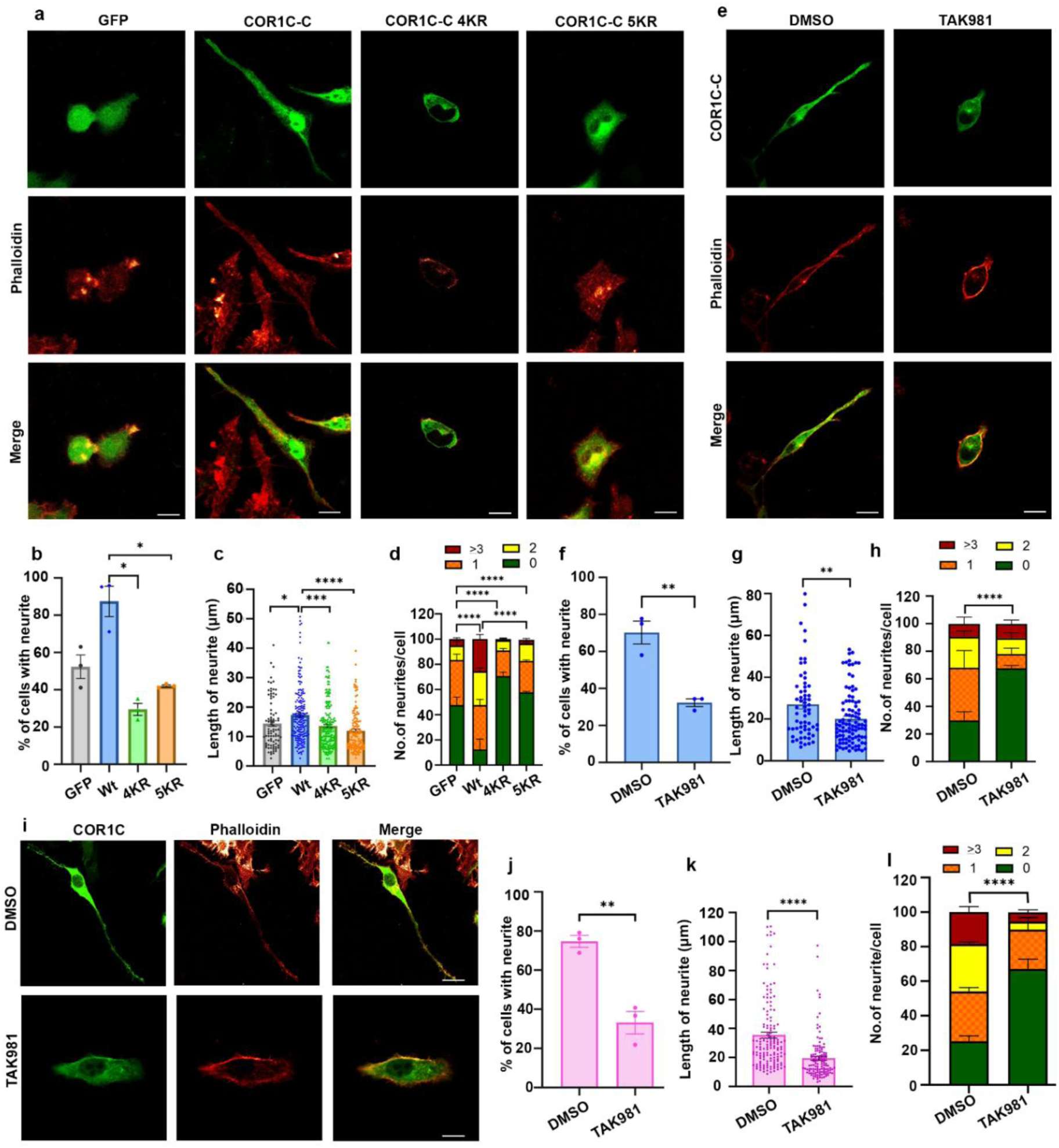
SUMOylation of COR1C promotes neurite formation. **a** Neuro2a cells were transfected with either GFP, GFP-tagged COR1C-C, COR1C-C-4KR, or COR1C-C-5KR and were imaged after 24 hours using a CLSM. The cells were stained with Phalloidin-647. Scale bar - 10 μm. **b** Percentage of cells with neurite(s) was quantified in 3 independent experiments (an average of 46 cells was counted each time). (*) indicate statistical significance with p-values of 0.0349 and 0.0348. **c** Average length of neurites per cell and **d** number of neurites of (a) were measured using ImageJ in 3 independent experiments and quantified; (*) and (****) indicate statistical significance with a p-value of 0.0189 and <0.0001, respectively. **e** CLSM imaging of GFP-COR1C-C expressing Neuro2a cells, treated with DMSO (control) or TAK981 (5 μM) for 48 hours. **f** Quantification of Neurite formation, **g** length of neurites and **h** number of neurites in (e). ∼ 50 cells were analyzed in 3 independent experiments. (**) indicate statistical significance with a p-value of 0.0044 and 0.0039, (****) indicate a p-value of <0.0001. **i** CLSM imaging of Neuro2a cells expressing GFP-COR1C, treated with DMSO (control) or TAK981 (5 μM) for 48 hours. **j** Quantification of Neurite formation and **k** length of neurites, and **l** number of neurites in (i). ∼ 50 cells were analyzed in 3 independent experiments. (**) and (****) indicate statistical significance with a p-value of 0.0013 and >0.0001, respectively. Statistical analysis was performed using an unpaired t-test for (b), (c), (f), (g), (j), and (k). Chi-square test was performed for (d), (h), and (l). Scale bar – 10 μm.

The importance of SUMOylation in neuritogenesis was further established by inhibiting global SUMOylation with TAK981. COR1C-C expressing N2A cells were treated with either DMSO or with TAK981, and the neurite forming ability of these cells was quantified. In COR1C-C expressing cells, TAK981 treatment resulted in a two-fold decrease (from 74.73% to 33.1%) in neurite formation as compared to DMSO control conditions (Fig. 5e, f). In addition, the neurite length in COR1C-C expressing cells also substantially decreased from 27±2.1 µm in DMSO-treated condition to 20±1.2 µm in TAK981-treated cells (Fig. 5g).

Further, we assessed whether diminished SUMOylation has an impact on the number of neurites formed per cell. The data suggest that there is no change in the percentage of cells with more than 3 neurites. But a careful glance at the quantification data revealed that an average of 51% of DMSO treated COR1C-C expressing cells have 1-2 neurites per cell, and this number decreases to 22% on average in TAK981-treated cells, highlighting the role of COR1C-C SUMOylation in neuritogenesis (Fig. 5h).

The effects of COR1C-C SUMOylation could be extrapolated to the COR1C full-length protein. Accordingly, when GFP-COR1C expressing cells were inhibited for SUMOylation by TAK981 treatment, similar decrease in the number of neurite positive cells (from an average of 68% to 43%), length of neurite (35±1.98 µm to 19.4±1.32 µm) and percentage of cells with 1-2 neurites per cell (from an average of 56% to 29.8%) were observed (Fig. 5i-l). From these observations, we conclude that the levels of COR1C and its SUMOylation are necessary for neurite formation, a hallmark of efficient differentiation of the neurons.

We further tested the importance of COR1C SUMOylation under conditions that promote neuronal differentiation. N2A cells were grown in a serum-deficient medium (1% FBS) for 72 hours, which induces neurite formation and differentiation (Fig. 6a). A larger fraction of the GFP control expressing cells presented a more rounded morphology, suggesting their undifferentiated state, and an average of 25% cells started showing signs of differentiation. In contrast, ∼ 64% of the serum-starved GFP-COR1C-C expressing N2A cells showed elongated neurites resembling axons and dendrites. Importantly, only 35.4% of GFP-COR1C-C-5KR expressing cells under serum-starved conditions could initiate differentiation (Fig. 6 b, c). This observation highlighted the importance of COR1C SUMOylation-dependent regulation of neurite formation and induction of neuronal differentiation.

**Fig. 6.**
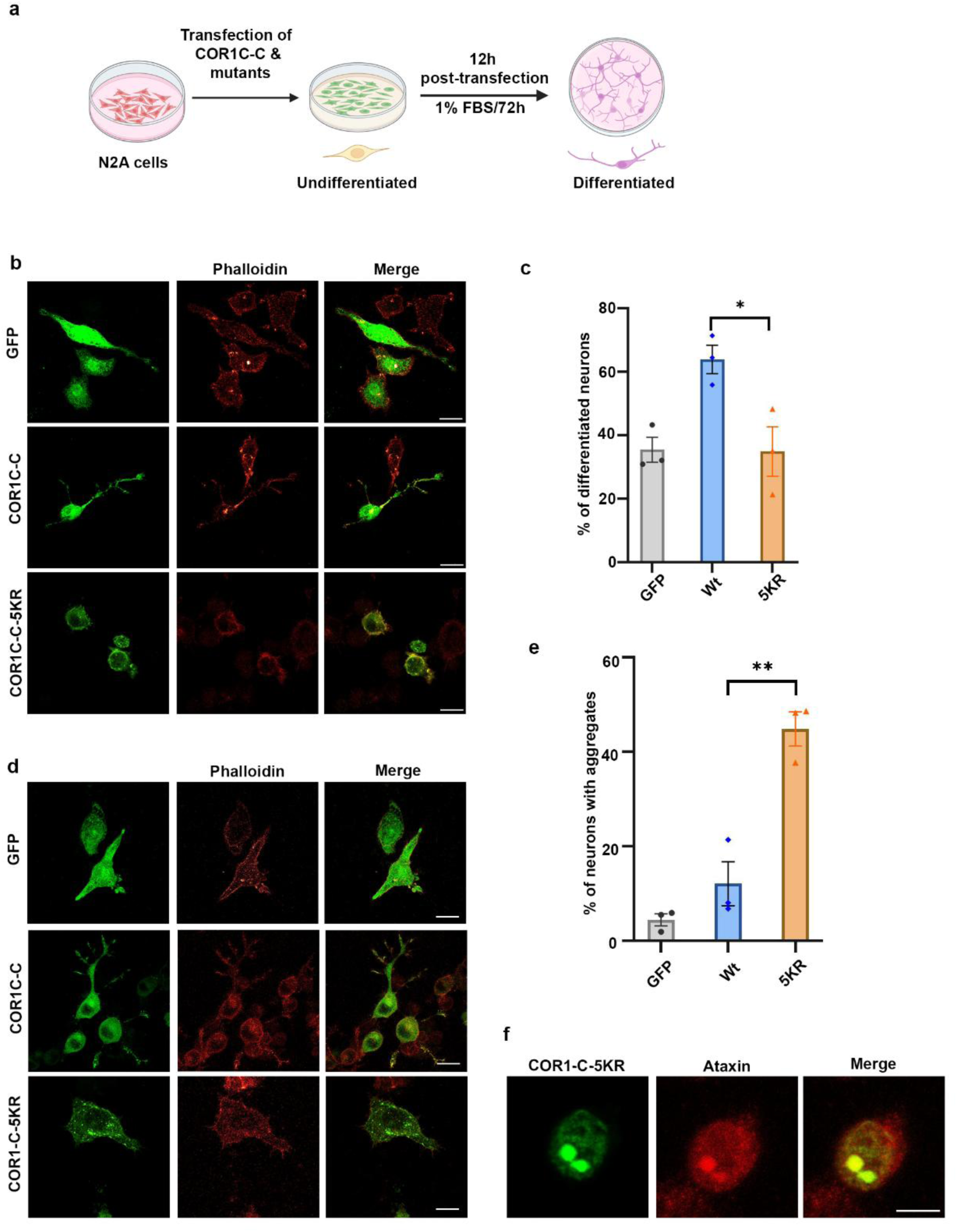
COR1C SUMOylation is crucial for the differentiation of Neuro2a cells.a. Detailed schematic illustration of Neuro2a differentiation under serum starvation. **b** Neuro2a cells were transfected with GFP, GFP-tagged COR1C-C, and COR1C-C-5KR and were imaged after 72 hours by using a CLSM. The cells were stained with Phalloidin-647. **c** Percentage of differentiated neurons was quantified in 3 independent experiments (an average of 45 cells was counted each time). (*) represents statistical significance with a p-value of 0.044. **d** Representative images of GFP, GFP-tagged COR1C-C, and COR1C-C-5KR expressing N2A cells for assessing cytoplasmic aggregates. **e** Percentage of neurons showing cytoplasmic aggregates was quantified in 3 independent experiments (∼ 50 cells were counted each time). (**) indicate statistical significance with a p-value of 0.0062. Statistical analysis was performed using an unpaired t-test. **f** Representative images of GFP-tagged COR1C-C-5KR expressing N2A cells for assessing cytoplasmic aggregates immunostained with anti-Ataxin2 antibody. Scale bar – 10 μm.

In an independent observation, we noticed that the GFP-COR1C-C-5KR mutant, when expressed under low serum conditions in N2A cells, forms cytosolic aggregates. These aggregates are significantly larger in number (∼ four-fold) when compared with GFP-COR1C-C wild-type expressing cells, suggesting a role for COR1C-C SUMOylation on its solubility status (Fig. 6d, e). Moreover, these aggregates observed for COR1C-C 5KR expression, when characterized for their nature using the stress granule marker Ataxin^35^, showed strong colocalization with Ataxin (Fig. 6f). This observation particularly suggested that COR1C SUMOylation can be a stress sensor and control its distribution in soluble versus aggregate forms when a particular stimulus, like differentiation, arrives.

Taken together, analyses of COR1C SUMOylation have suggested that SUMOylation of COR1C plays a critical role in actin-filament dynamics required for the number and size of cellular projections, such as filopodia and neurites. Additionally, this SUMOylation is required to maintain COR1C solubility and aggregation properties during differentiation (Fig. 7).

**Fig. 7.**
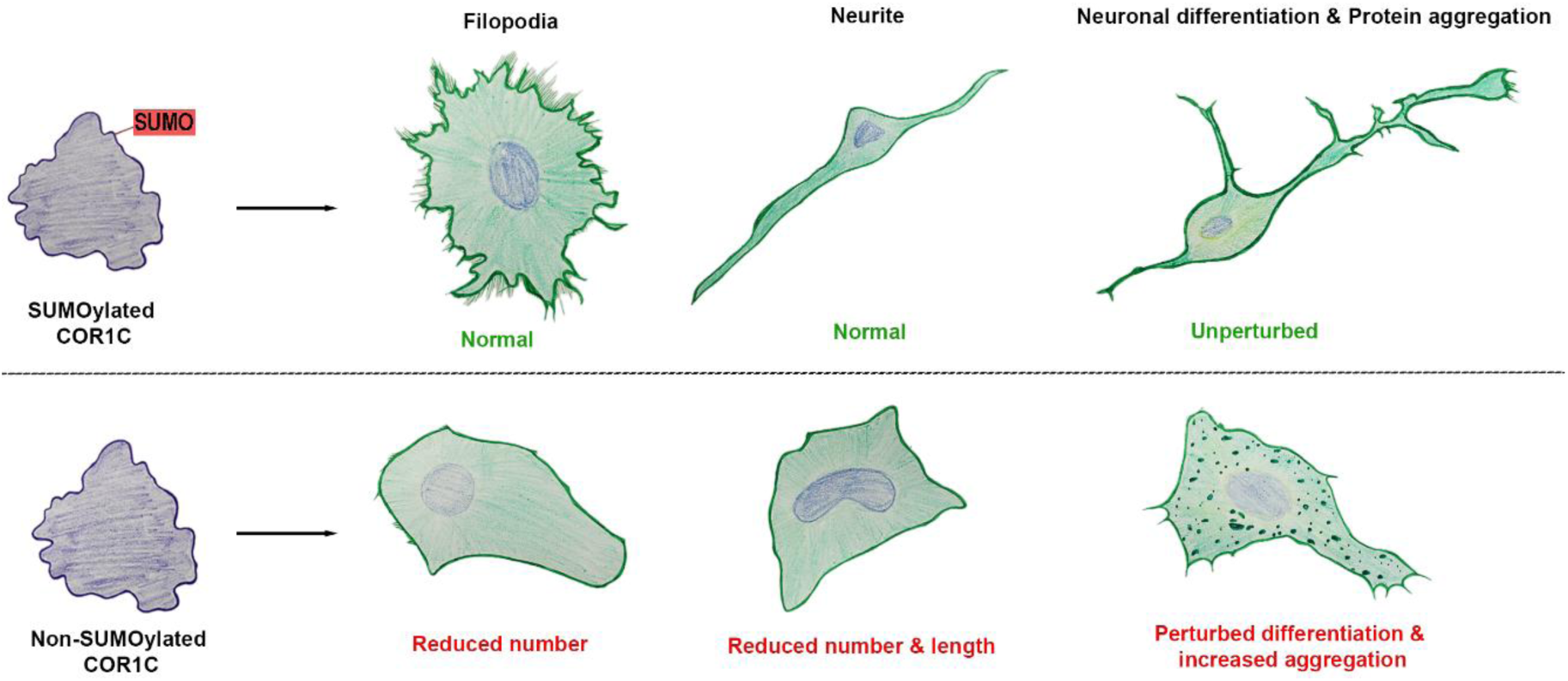
SUMOylation of Coronin1C is critical for filopodia formation and neuronal differentiation. F-actin binding protein COR1C is modified by SUMO1 in its carboxy-terminus. SUMOylated COR1C promotes the assembly of actin-rich structures like filopodia and neurites. Non-SUMOylatable COR1C impairs assembly of these actin-rich structures, causing a decrease in filopodial numbers to impede cellular migration and stunted neuronal differentiation through neuritogenesis. SUMOylation of COR1C is important for neuronal differentiation and maintenance of the COR1C soluble pool. Accordingly, non-SUMOylatable COR1C forms neuronal aggregates, a hallmark of neurodegenerative diseases (NDDs), even in the absence of any cytoskeletal or physiological stress.

## Discussion

In recent years, SUMOylation of Periplakin, Cofilin, Vimentin, intermediate filaments, and actin microfilament-binding proteins has been characterized in detail, and they regulate cytoskeleton dynamics and stability^23,37^.

In this context, we report SUMOylation of an actin-binding protein, COR1C. The carboxy-terminal region of COR1C, important for actin interaction, is SUMOylated at five distinct lysine residues by SUMO1 (Fig. 1, 2). We observed that SUMOylation of COR1C plays a crucial role in filopodia formation, indicated by a reduction in the number of filopodial projections in epithelial cells (Fig. 3). COR1C has been previously associated with lamellipodia formation by trafficking Rac1 to the leading edge of the cell^27^. It regulates actin dynamics at the leading edge of the cell. Our observation that COR1C SUMOylation likely occurs at K329, K403, K413, K440, and K464 residues lying in the proximity of actin-binding residues K418, K419, K427, and K428 of COR1C^14^ can affect not only the binding of COR1C with actin, but also with other actin-binding proteins. This, in turn, regulates actin dynamics and contributes to the formation of filopodial cellular projections. Cdc42, a small GTPase required for actin filament dynamics, affects filopodia formation^31^. The filopodia numbers do not reduce drastically in COR1C-expressing and ML141-treated cells, suggesting COR1C-C works synergistically with Cdc42. Further, a drastic decrease in the filopodial numbers in the ML141 treated cells expressing the COR1C-C-5KR highlights the importance of COR1C-C SUMOylation in regulating actin dynamics and filopodia formation. Even though evidence for a direct interaction between Cdc42 and COR1C is unavailable, a strong indication of the possibility of their direct or indirect association at the cellular edges by actin-regulating proteins is undeniable. Moreover, a recent study established COR1C SUMOylation-dependent interaction with ARP2 to orchestrate actin dynamics in lung adenocarcinoma^38^. The correlation of COR1C SUMOylation, Cdc42, and ARP2 can be further explored to better understand the mechanism underlying filopodia and neurite formation. It is important to note that coordinated COR1C and ARP2/3 complex activity is required for actin filament branching, which is essential for the formation of filopodia-like structures. Several previous studies have established that COR1C is important for cellular migration^10^. Our results indicate that SUMOylation of COR1C is required for the formation of filopodia-like structures at the leading edge of a cell for controlling cellular migration.

Neurites are filopodia-derived structures rich in actin and tubulin cytoskeletal networks. Neurites are of utmost importance during neuronal differentiation as neurites further differentiate to mature into axons and dendrites of a neuron^33^. Involvement of COR1C in neuronal morphology and migration, in addition to COR1C localization at the base of a neurite, has been reported with an emphasis on the role of the carboxy-terminal region (COR1C-C) in neurite formation^13^. Thus, a decrease in neurite formation, its length, and numbers per cell in non-SUMOylatable COR1C-C expressing N2A cells places a focus on PTM-based regulation of cytoskeletal dynamics during neurite formation (Fig. 4). Our study highlights the importance of COR1C SUMOylation in the formation of actin-rich neurites, with a regulatory role assigned to the SUMOylation of COR1C-C during neural differentiation.

It is evident that neuronal health can be altered by protein aggregation, which is one of the key events observed in multiple neurodegenerative diseases (NDDs)^39^. Several of these NDD-linked proteins, like tau, synuclein, etc., are cytoskeletal network proteins, and they tend to form aggregates in the neurons in the non-SUMOylated state^40^. The non-SUMOylatable COR1C-C-5KR also forms visible aggregates in N2A cells; however, only under growth-restricting and differentiation-favouring low-serum stress conditions (Fig. 5). The same protein fails to assemble into aggregates under normal growth conditions. This may imply that SUMOylation is critical for maintaining a soluble pool of COR1C. The relative abundance of the aggregate pool is inaccessible and possibly incapable of modulating actin dynamics at cellular protrusions. SUMOylation of COR1C helps maintain its soluble pool in the cytoplasm. It remains to be seen if low serum-induced growth restriction can serve as a stress that COR1C SUMOylation can help manage.

SUMOylation of a target protein can impact its protein-protein interactions. COR1C interacts with multiple proteins other than actin, like Rac1^27^, Cofilin^41^, and ARP2/3^18^. Each of these proteins is known to affect actin filament dynamics. It will be of interest to analyze the impact of SUMOylation on COR1C interactions. It will also help delineate the exact mechanism by which COR1C SUMOylation regulates actin remodelling in cellular protrusions. Our observations also argue in favour of understanding the consequences of polypeptide-based PTMs like SUMOylation in cytoskeletal dynamics contributing to cellular growth, migration, and differentiation.

## Materials and methods

### Plasmid constructs and antibodies

Coronin1C was amplified with appropriate primers from HeLa cell cDNA. The following specific primers were used: COR1C Forward: 5’-ATAGAATTCATGAGGCGAGTGGTA-3’; COR1C Reverse: 5’-TATGGTCGACTCGGCTGCTATCTTTGC-3’. COR1C full-length and C-term (aa 315-474) constructs were cloned into vectors pEGFP-C2 for mammalian cell expression and visualization analyses or into pGEX-6P-1 for protein purification and SUMOylation analyses. The COR1C lysine mutants were generated using the Q5 Site-Directed Mutagenesis Kit (NEB) according to the manufacturer’s protocol. pSUMO1/2 was a gift from Primo Schaer (Department of Biomedicine, University of Basel; Addgene plasmid # 52258, #52259)^42^. The following antibodies were used for Western blotting: anti-GST (1:5000, HiMedia-HTI009), anti-His (1:2000, #BB-AB0010S, Biobharti), anti-SUMO1 (76-86, DSHB) anti-GFP (1:5000; #sc-9996 Santa Cruz Biotechnology), anti-GAPDH (1:3000; #10011 Abgenex), anti-COR1C (Proteintech-14749-1-AP), anti-Flag (Sigma, F1804), Anti-Ataxin2 (Proteintech-21776-1-AP), Goat anti-mouse AlexaFluor Plus 800 (1:40,000, #A32730, Thermo Fisher Scientific), Goat anti-rabbit AlexaFluor Plus 680 (1:40,000, #A32734 Thermo Fisher Scientific). Phalloidin-647 (1:100, PHDN1-A, Cytoskeleton Inc.).

### *In bacto* SUMOylation

Coronin1C constructs cloned in the pGEX-6P-1 vector were co-transformed into *E. coli* with pSUMO1 or pSUMO2 plasmids. Double transformants were selected on Luria Bertani (LB) agar plates having 50 mg/L of ampicillin (Amresco, #0339) and 25 mg/L of streptomycin (SRL #19014). For *in bacto* SUMOylation, mid–log phase *E. coli* cultures were induced with 200 μM isopropyl β-D-1-thiogalacto-pyranoside (IPTG) (HiMedia #RM2578) at 30 °C for 4 h. The protein was purified according to the detailed protocol described elsewhere^37,43^. Cells were harvested by centrifugation, and soluble protein fractions were extracted by sonication in a buffer containing 20 mM Tris, pH 7.5, 150 mM NaCl, 1 mM EDTA, 0.1% Triton, 5 mM β-mercaptoethanol (MP biomedicals #194705), and 1 mM phenylmethylsulfonyl fluoride (MP Biomedicals #195381). Crude lysates were then cleared by centrifugation at 20,000 × g at 4 °C for 30 min. Proteins were affinity-purified on Glutathione beads (Merck #70541), followed by three washes with the same buffer containing 400 mM NaCl. Proteins bound to beads were extracted by boiling in the Laemmli buffer and subsequently analyzed on SDS–PAGE/Western blotting. To cleave GST from the proteins, the bead-bound proteins were incubated with PreScission protease in a 1:50 ratio for 12 h at 4 °C. For deconjugation experiments, bead-bound protein fractions were treated with SENP1 catalytic fragment in 1:50 ratio at room temperature for 1 h and analyzed on SDS-PAGE/Western blotting.

### Cell culture and transfections

HEK293T (ATCC no. CRL-3216), HeLa (ATCC no. CCL2), MCF7 (ATCC no. HTB-22), and Neuro2a (ATCC no. CCL131) cell lines were mycoplasma free, and were cultured in DMEM (Gibco) supplemented with 10% (v/v) fetal bovine serum (Gibco) and 1% antibiotics (Gibco) in a humidified incubator at 37 °C under 5% CO_2_ conditions. The cells were grown in 100-mm cell culture plates (1 × 10^7^ cells) or in six-well plates (3 × 10^5^ cells per well) for Western blot analysis. After 12 h of plating, cells were transfected using polyethylenimine (PEI) 25-kDa linear (23966-2g; Polysciences Corporation Ltd.) as previously described^34^. To induce neuronal differentiation, Neuro2a cells were grown in low serum (1% FBS) for 12 h post-transfection and allowed to grow for 72 h. For SUMOylation inhibition, 8-12 h post-transfection, transfected cells were treated with 5 μM TAK981 (MedChemExpress #HY-111789) for the suggested durations and processed further. For Cdc42 inhibition, transfected cells were treated with 10 μM ML141 (CAS No. 71203-35-5, ADSci #AD201975) for 2 h under regular growth media at 37 °C and processed further.

### *In cellulo* SUMOylation assay

HEK293T cells were co-transfected with Venus-SUMO1 and pEGFP-Coronin1C. 48 hours post-transfection, the total cell lysates were prepared in RIPA buffer (50 mM Tris, 10 mM EDTA, 0.1% Sodium deoxycholate, 1% Triton X-100, 1 mM EGTA, 0.1 % SDS) supplemented with 40 mM N-ethyl maleimide (NEM, Sigma-04259). The lysate was sonicated and centrifuged, and the total soluble protein was boiled in 1X Laemmli buffer for further analysis.

For Immunoprecipitation, HEK293T cells were co-transfected with pCMV-3tag-1a-COR1C-C constructs along with pEGFP-SUMO1G/GG or SUMO2G/GG (a kind gift from Dr. Jomon Joseph, NCCS, Pune) in a 2:1 ratio using PEI. 48 h post-transfection, cells were lysed in lysis buffer (50 mM Tris, 10 mM EDTA, 0.1% Sodium deoxycholate, 1% Triton X-100, 1 mM EGTA, 0.1 % SDS) supplemented with 40 mM NEM, PMSF, and protease inhibitor cocktail. Lysed cells were sonicated at 35% amplitude with 3 sec pulses for 1 minute, followed by centrifugation at 14000 rpm for 30 minutes at 4 °C. Total supernatant was bound to 15 μL anti-Flag M2 affinity gel (Merck-A2220) and incubated for 16 h at 4 °C with constant rotation. The protein-bound beads were washed three times with PBS and boiled in Laemmli buffer for Western blotting.

### Western blotting

Proteins were separated on SDS–PAGE and then transferred to polyvinylidene difluoride (PVDF) membrane. The membranes were incubated in blocking buffer, 5% bovine serum albumin in TBS with 0.1 % Tween-20 (TBS-T), for 1 h at room temperature. After incubation with the indicated primary antibodies, membranes were given three washes of 5 min each with TBS-T, followed by incubation with respective AlexaFluor-conjugated secondary antibodies for 1 h at room temperature. Detection of protein bands was carried out using the Li-COR IR system (Model: 9120).

### Immunofluorescence microscopy

HeLa, MCF7, and Neuro2a cells grown on glass coverslips were transfected with the indicated constructs. Cells were fixed with 4% formaldehyde (Merck-1.94900) for 5 min, followed by 10 min of incubation in rehydration buffer (PBS, 0.1% Triton). Cells were then incubated with Phalloidin-647 diluted 1:100 in PBS. Cells were washed thrice with PBS, and coverslips were mounted on glass slides using Fluoroshield mounting medium (Sigma-F6057). Confocal imaging was performed using an Olympus Confocal Laser Scanning Microscope (CLSM)–FV3000.

### Wound healing assay

HeLa cells were grown to 75% confluency in a 6-well plate or 3 cm dish. It was then transfected with the desired plasmids. The media was changed after ∼ 12 h. The cells were allowed to grow until ∼ 95% confluency was reached. Scratches were made gently using a 2 μL microtip on the surface of the dish by holding the tip vertically without tilting. After making the scratches, the dish was washed gently with PBS to remove the debris without disturbing the cells. The cells were now grown in DMEM with 1 % FBS and imaged at required time points using an inverted microscope (Leica Microsystems Model-DMIL LED Fluo). The wound closure rate was quantified using TScratch software.

### Statistics and Reproducibility

Image analysis, processing, and filopodia quantification were performed using ImageJ software. Band intensities were quantified using GelQuant.NET. The experiments were independently repeated at least three times, and the values are expressed as mean ± SEM. The p-values were calculated using Student’s t-test; p ≤ 0.05 was considered statistically significant. Graphs were plotted using GraphPad Prism, and figures were arranged in Adobe Photoshop 3.0.

## Supporting information

Supplementary_COR1C_MS

## Acknowledgements

The authors acknowledge the efforts of Shaunak Vilas Joshi for his help with clone generation. The authors are thankful to Jyotsna Kawadkar for her help in arranging the figures. We are grateful to Dr. Dhananjay Huilgol and Mr. Sumit Singh for their efforts in performing key experiments at BRIC-NBRC, Manesar. G.J. is supported with a fellowship by IISER Bhopal. HVS is supported by DBT for a fellowship. This work was supported by research grants CRG/2020/000496 from SERB, IIRPSG-2024-01-01766 from ICMR, and intramural support from IISER Bhopal to RKM.

## Author contributions

G.J. designed and performed all experiments and analyzed and interpreted data. H.V.S. helped with the *in bacto* experiments. R.K.M. conceptualized the project and analyzed and interpreted data. G.J. and R.K.M. wrote the manuscript, and R.K.M. obtained the funding.

## Supplementary information

**Fig. S1.**
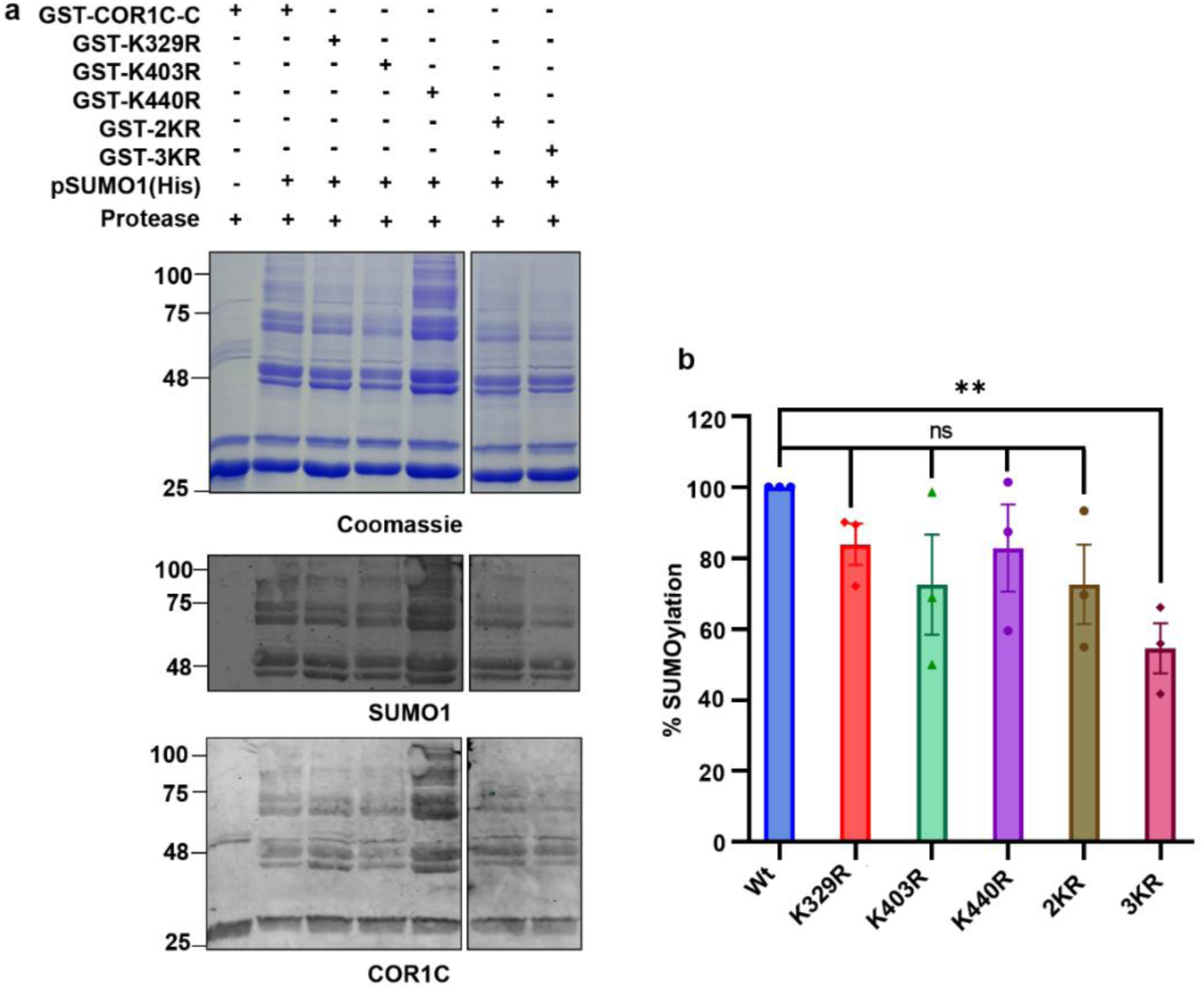
COR1C is SUMOylated at multiple lysines in its C-terminus. SDS-PAGE analysis of *in bacto* SUMOylation reaction with GST-COR1C-C wild-type and mutant post digestion with PreScission protease to produce untagged COR1C-C (modified and unmodified, upper panel). Immunoblotting of samples as upper panels with anti-SUMO1 (middle panel) and anti-COR1C (lower panel) antibodies.

**Fig. S2.**
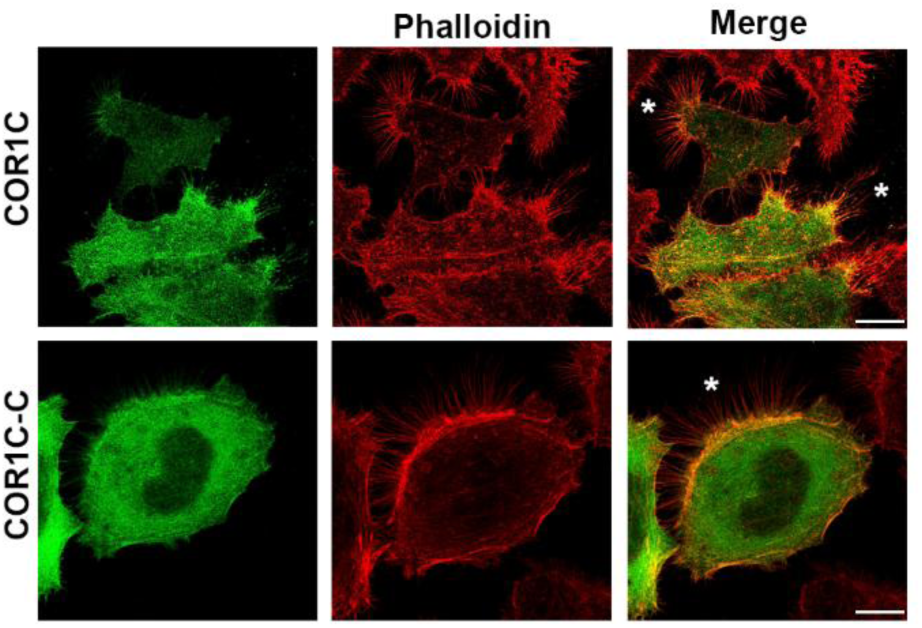
The carboxy-terminus of Coronin1C localizes at the filopodial projections. HeLa cells transfected with GFP-COR1C and GFP-COR1C-C stained with Phalloidin-647. Scale bar - 10 µm. White stars denote the filopodial projections.

**Fig. S3.**
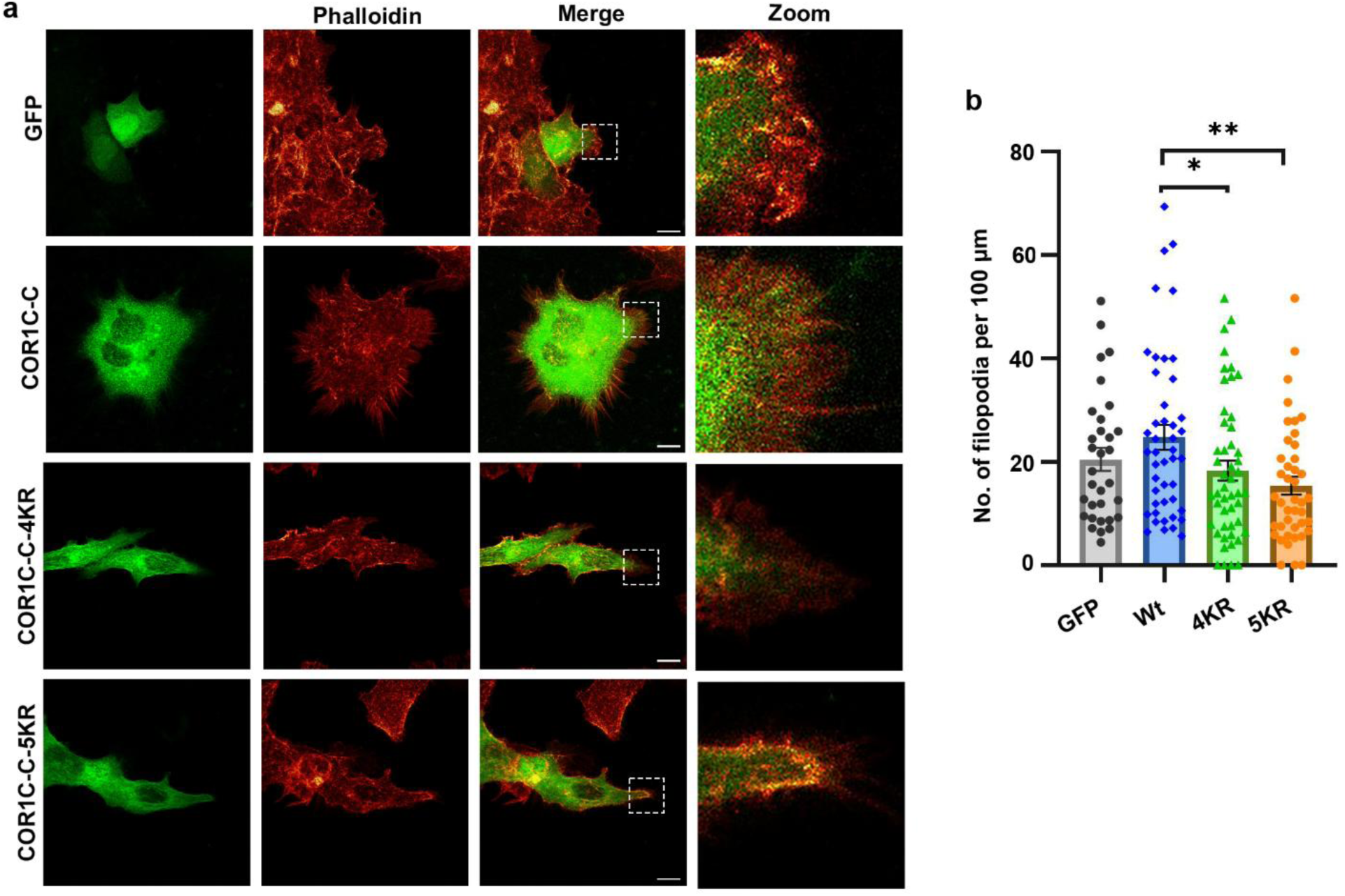
COR1C-C SUMOylation is critical for filopodia formation in cells. **a** CLSM imaging of MCF7 cells transfected either with GFP-tagged COR1C-C, COR1C-C-4KR, or COR1C-C-5KR. Cells were stained with Phalloidin-647. Dotted-box highlights filopodial structures in the leading edges, as shown in the zoom. **b** Filopodial projections as seen in (a) were quantified using the FiloQuant plugin of ImageJ (∼ 50 cells were analyzed in three independent experiments). Statistical analysis was performed using an unpaired t-test; (*) and (**) indicate statistical significance with a p value of 0.0389 and 0.0026. Scale bar - 10 µm.

**Fig. S4.**
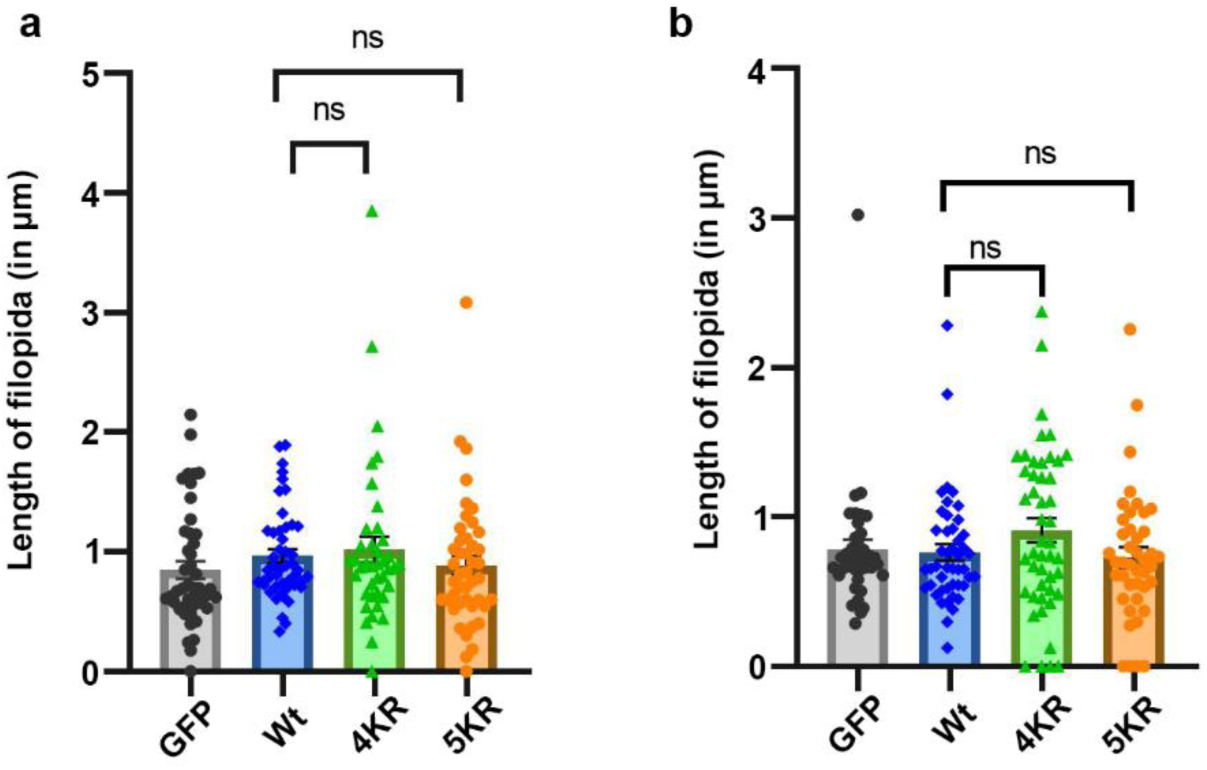
COR1C-C SUMOylation does not affect filopodia length in cells. Quantification of the length of filopodial projections measured from HeLa cells **(a)** and MCF7 cells **(b)**, as shown in Figure S3, was performed using the FiloQuant plugin of ImageJ (∼ 50 cells were analyzed in three independent experiments). Statistical analysis was performed using an unpaired t-test.

**Fig. S5.**
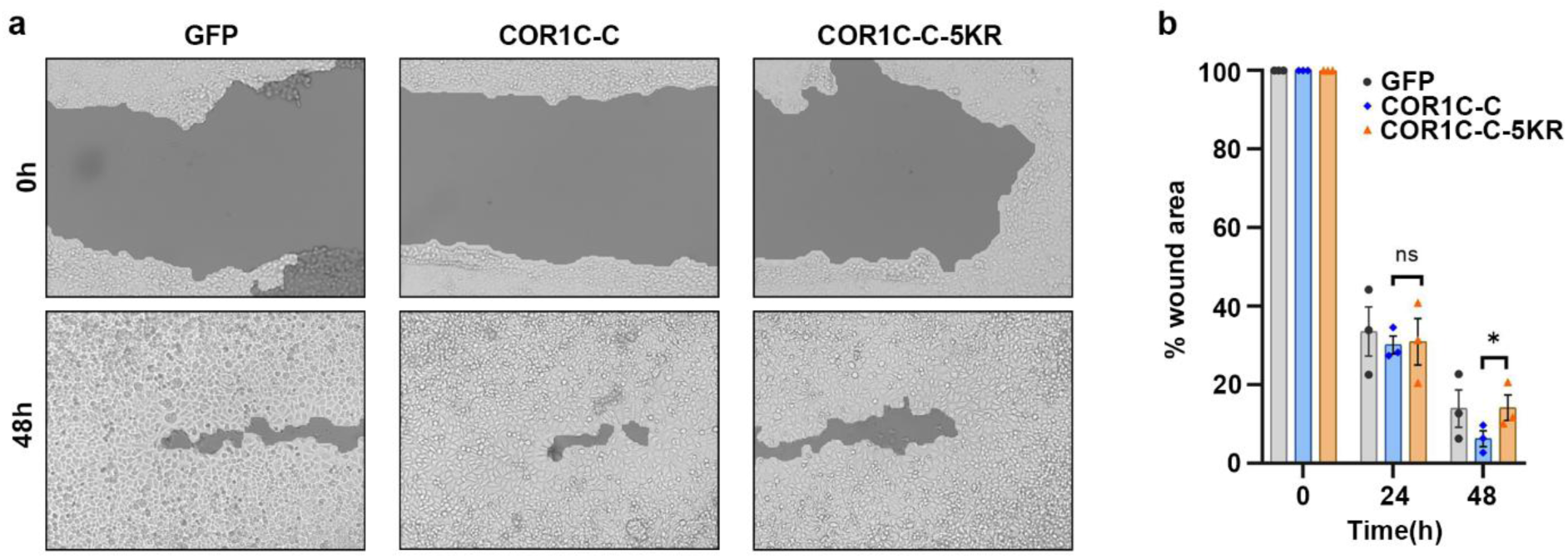
COR1C SUMOylation is required for efficient cellular migration. **a** Representative image of the wound healing assay performed with HeLa cells expressing GFP, GFP-COR1C-C, or GFP-COR1C-C-5KR. Images were captured at 10x magnification using an inverted microscope. **b** Quantification of wound area closure using TScratch software. Images are representative of n=3 experiments. Data was analysed by using a paired t-test, and (*) indicates statistical significance with a p-value of 0.0398.

**Table S1.**
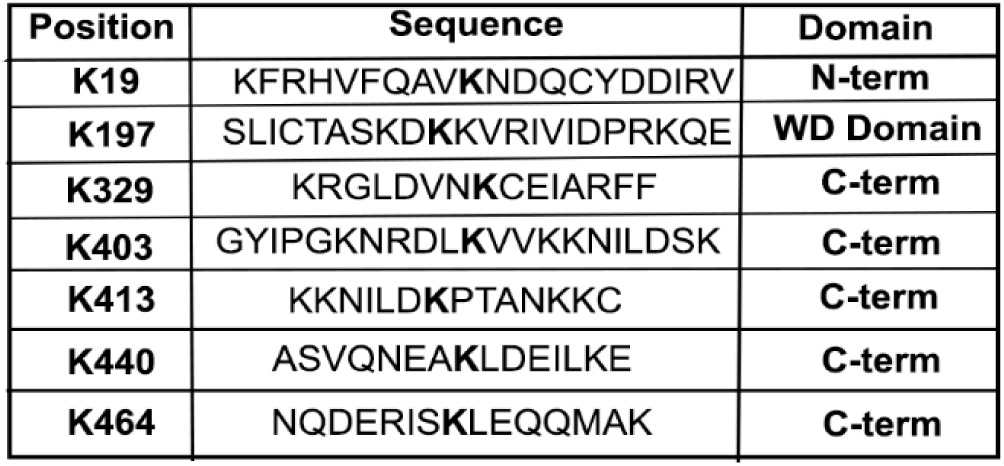
*in silico* prediction of COR1C SUMO sites.

