## Supplementary_COR1C_MS for "Coronin1C SUMOylation modulates filopodia formation, neuritogenesis, and neuronal differentiation"

### Supplementary information

| Position | Sequence | Domain |
|---|---|---|
| **K19** | KFRHVFQAV**K**NDQCYDDIRV | **N-term** |
| **K197** | SLICTASKD**K**KVRIVIDPRKQE | **WD Domain** |
| **K329** | KRGLDVN**K**CEIARFF | **C-term** |
| **K403** | GYIPGKNRDL**K**VVKKNILDSK | **C-term** |
| **K413** | KKNILD**K**PTANKKC | **C-term** |
| **K440** | ASVQNEA**K**LDEILKE | **C-term** |
| **K464** | NQDERIS**K**LEQQMAK | **C-term** |

**Table S1 | *in silico* prediction of COR1C SUMO sites**

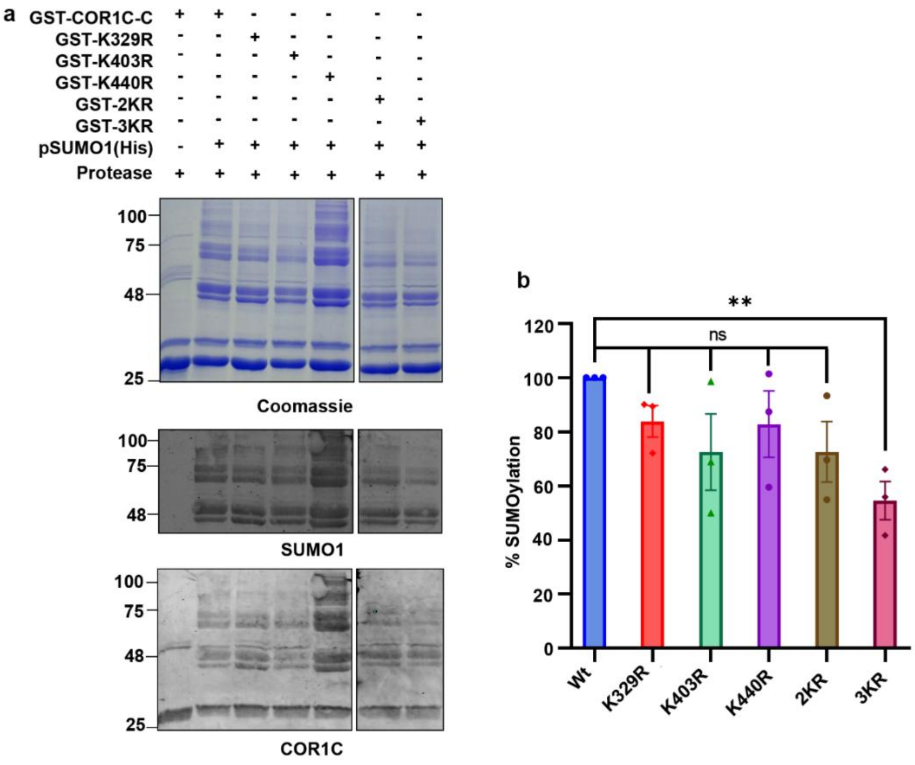


**Fig. S1 | COR1C is SUMOylated at multiple lysines in its C-terminus.**

SDS-PAGE analysis of *in bacto* SUMOylation reaction with GST-COR1C-C wild-type and mutant post digestion with PreScission protease to produce untagged COR1C-C (modified and unmodified, upper panel). Immunoblotting of samples as upper panels with anti-SUMO1 (middle panel) and anti-COR1C (lower panel) antibodies.

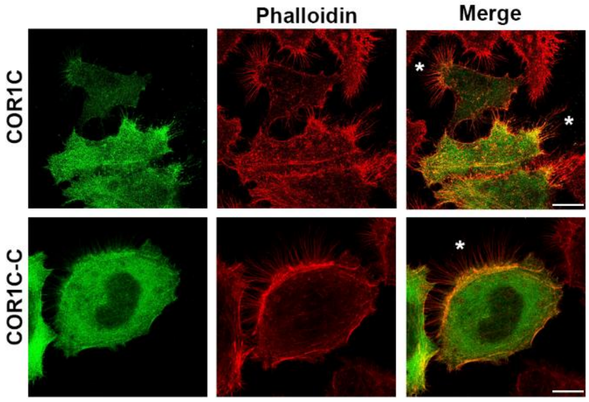


**Fig. S2 | The carboxy-terminus of Coronin1C localizes at the filopodial projections**.

HeLa cells transfected with GFP-COR1C and GFP-COR1C-C stained with Phalloidin-647.

Scale bar - 10 μm. White stars denote the filopodial projections.

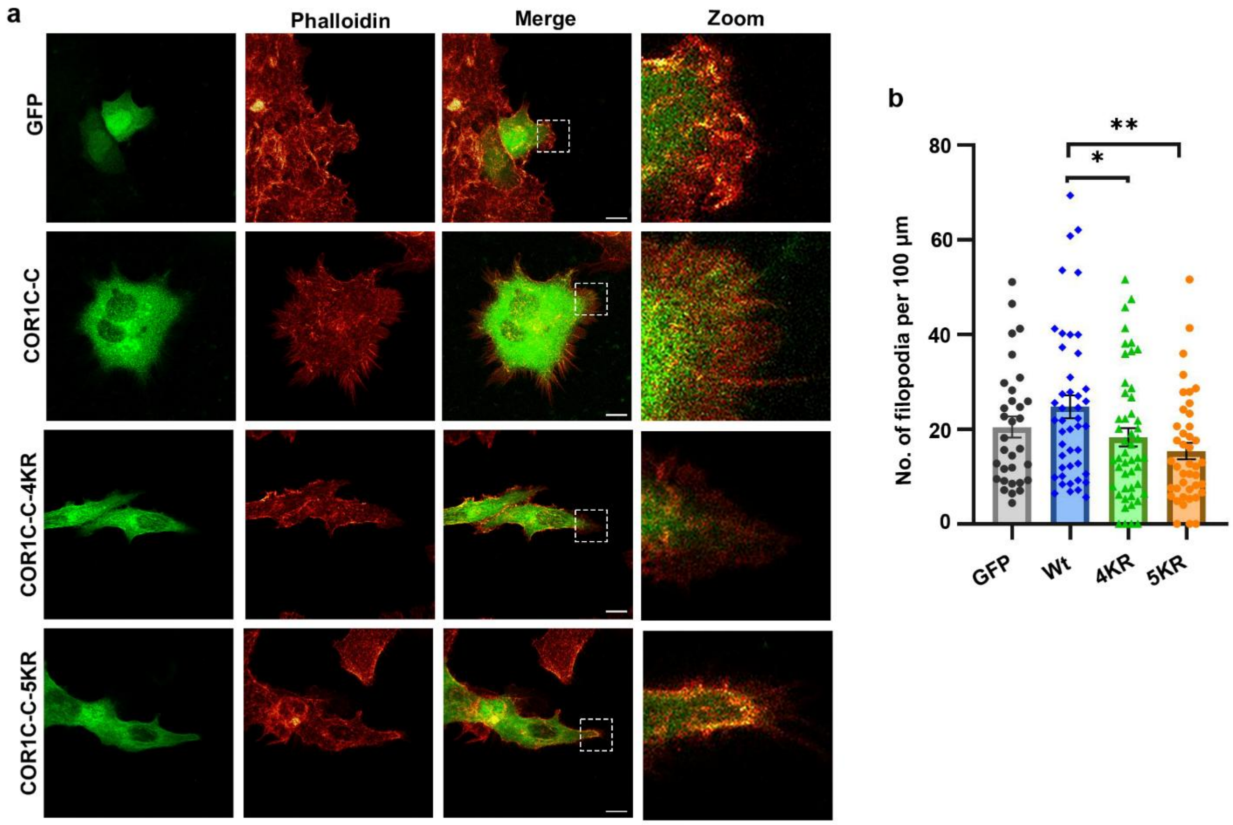


**Fig. S3 | COR1C-C SUMOylation is critical for filopodia formation in cells.**

**a** CLSM imaging of MCF7 cells transfected either with GFP-tagged COR1C-C, COR1C-C-4KR, or COR1C-C-5KR. Cells were stained with Phalloidin-647. Dotted-box highlights filopodial structures in the leading edges, as shown in the zoom. **b** Filopodial projections as seen in (a) were quantified using the FiloQuant plugin of ImageJ (~ 50 cells were analyzed in three independent experiments). Statistical analysis was performed using an unpaired t-test; (*) and (**) indicate statistical significance with a p value of 0.0389 and 0.0026. Scale bar - 10 µm.

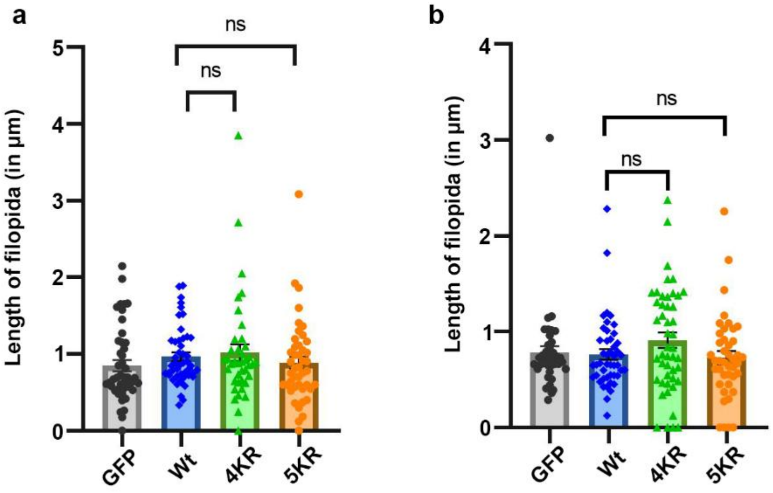


**Fig. S4 | COR1C-C SUMOylation does not affect filopodia length in cells.**

Quantification of the length of filopodial projections measured from HeLa cells **(a)** and MCF7 cells **(b)**, as shown in Figure S3, was performed using the FiloQuant plugin of ImageJ (~ 50 cells were analyzed in three independent experiments). Statistical analysis was performed using an unpaired t-test.

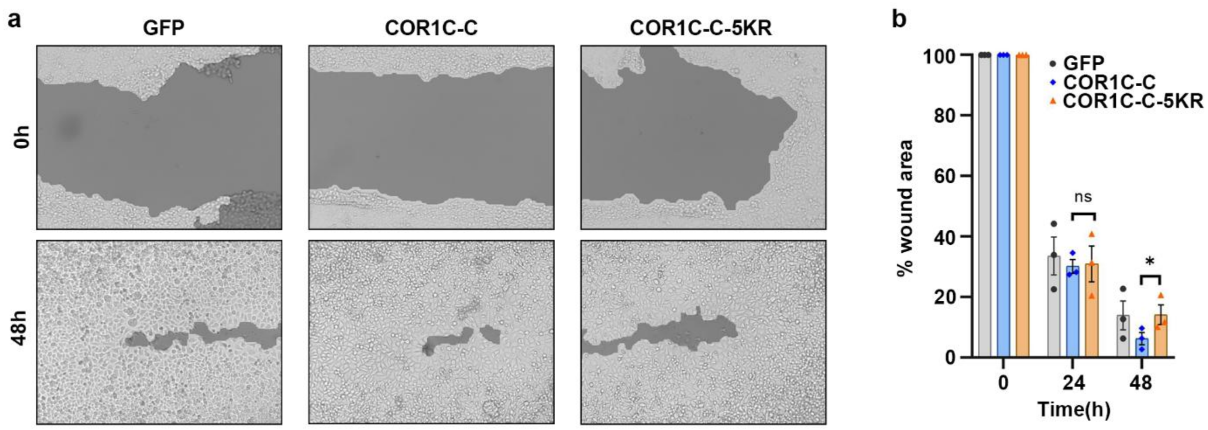


**Fig. S5 | COR1C SUMOylation is required for efficient cellular migration.**

**a** Representative image of the wound healing assay performed with HeLa cells expressing GFP, GFP-COR1C-C, or GFP-COR1C-C-5KR. Images were captured at 10x magnification using an inverted microscope. **b** Quantification of wound area closure using TScratch software. Images are representative of n=3 experiments. Data was analysed by using a paired t-test, and (*) indicates statistical significance with a p-value of 0.0398.
